# Uncoupling Intraflagellar Transport and Primary Cilia Formation Demonstrates Deep Integration of IFT in Hedgehog Signaling

**DOI:** 10.1101/226324

**Authors:** Thibaut Eguether, Fabrice P Cordelieres, Gregory J Pazour

## Abstract

The vertebrate hedgehog pathway is organized in primary cilia and hedgehog components relocate into or out of cilia during signaling. Defects in intraflagellar transport (IFT) typically disrupt ciliary assembly and attenuate hedgehog signaling. Determining if IFT drives the movement of hedgehog components is difficult due to the requirement of IFT for building cilia. Unlike most IFT proteins, IFT27 is dispensable for cilia formation but affects hedgehog signaling similar to other IFTs allowing us to examine its role in the dynamics of signaling. Activating signaling at points along the pathway in *Ift27* mutant cells showed that IFT is extensively involved in the pathway. Similar analysis of *Bbs* mutant cells showed that BBS proteins participate at many levels of signaling but are not needed to concentrate Gli transcription factors at the ciliary tip. Our analysis showed that smoothened delivery to cilia does not require IFT27, but the role of other IFTs is not known. Using a rapamycin-induced dimerization system to stop IFT after ciliary assembly was complete we show that smoothened delivery to cilia is IFT independent.

**Abbreviations:**

MEFs: mouse embryonic fibroblasts
SAG: smoothen agonist
IFT: intraflagellar transport
FKBP: FK506 Binding Protein 12
FRB: FKBP12-rapamycin binding

## Introduction

Primary cilia are microtubule-based hair-like sensory organelles that are crucial regulators of cell signaling. The receptors and downstream effectors of many signaling proteins are enriched and sequestered in the cilium and this compartmentalization allows for fine temporal and spatial regulation of pathway activation as well as downstream signal propagation. In vertebrates, the best studied cilia-linked pathway is sonic hedgehog, which plays fundamental roles during development and in adult tissue homeostasis. All the key components of the pathway are enriched in the cilium (Corbit *et al*, 2005; Haycraft *et al*, 2005; Ocbina & Anderson, 2008; Rohatgi & Scott, 2007) and their localization changes dynamically in response to the activation of the pathway. In its off-state, patched-1 (Ptch1), the hedgehog ligand receptor, accumulates in the primary cilium and prevents ciliary accumulation and the activation of smoothened (Smo). Upon binding of ligand, Ptch1 is removed from the cilium, Smo is derepressed and accumulates in the cilium. Smo subsequently activates downstream signaling, which results in the accumulation of the Gli family of transcription factors at the tip of the cilium before their modification and translocation to the nucleus where they modulate target genes.

The movement of hedgehog pathway components has been well studied. Ptch1 and Smo come and go from the cilium depending on the activation status of the pathway (Corbit *et al*, 2005; Rohatgi & Scott, 2007; Kim *et al*, 2009). Gli2 and Gli3 localize to the tip of primary cilia where they accumulate to high levels upon pathway activation (Haycraft *et al*, 2005; Kim *et al*, 2009; Keady *et al*, 2012; Santos & Reiter, 2014). SuFu localizes to the primary cilium in a Gli-dependent manner (Haycraft *et al*, 2005; Zeng *et al*, 2010; Tukachinsky *et al*, 2010). Kif7, like the Gli factors, localizes at the ciliary tip and accumulates when the pathway is activated (He *et al*, 2014). However, mechanisms underlying their trafficking to and within the cilium remain elusive. Part of these movements is facilitated by intraflagellar transport (IFT) (Eguether *et al*, 2014; Keady *et al*, 2012) and perturbing IFT disrupts hedgehog signaling (Huangfu *et al*, 2003; Liem *et al*, 2012; Duran *et al*, 2016). However, it is difficult to know whether IFT is participating directly in the transport of many of the components as perturbing IFT typically disrupts ciliary assembly which could cause indirect effects. Hence, it is fundamental to experimentally disconnect IFT and primary cilia formation to better understand the function of the IFT system in hedgehog signaling. Previously, we showed that the complex B proteins - IFT25 and IFT27 - are dispensable for cilia formation but are fundamental for the regulation of hedgehog signaling (Keady *et al*, 2012; Eguether *et al*, 2014), thus describing a system in which IFT and primary cilia formation are naturally uncoupled. In this study, we further examined the role of IFT27 in hedgehog signaling and compared it to the BBSome. We find that IFT27 is involved in most of the major steps of hedgehog signal transduction and unexpectedly plays roles independent of the BBSome.

Our work indicates that IFT25 and IFT27 are extensively involved in hedgehog signaling, but the question of whether other IFT proteins have direct roles in signaling remains. To test this, we designed a system to perturb IFT in fully formed cilia and asked how this affects hedgehog signaling. Our approach uses an in-cell dimerization system based on rapamycin proprieties. Rapamycin can simultaneously bind the FK506 Binding Protein 12 (FKBP) and the FKBP12-rapamycin binding (FRB) domain of mTOR with high affinity (Chen *et al*, 1995). Proteins that would not normally dimerize can be tagged with the FKBP and FRB domains and be brought together upon addition of rapamycin (Bayle *et al*, 2006). The system is highly efficient and has been used to study receptor-mediated endocytosis (Varnai *et al*, 2006), clathrin-coated vesicles (Robinson *et al*, 2010), actin network (Castellano *et al*, 1999) and the gating properties of the transition zone of primary cilia (Lin *et al*, 2013). In our system, an IFT protein is tagged with the FKBP domain while the FRB domain is sequestered at the mitochondria. In the absence of rapamycin, the IFT-FKBP fusion protein participates in IFT and assembles cilia. Upon addition of rapamycin The IFT is sequestered at the mitochondria and no longer supports IFT.

## Results

### IFT27 is necessary to propagate hedgehog signaling downstream of smoothened

*Ift27* mutants fail to remove Ptch1 from cilia upon activation of hedgehog signaling and fail to keep Smo from accumulating in cilia of non-activated cells indicating that IFT27 acts in the very early steps of pathway activation (Eguether *et al*, 2014). To understand if IFT27 is also acting downstream of Smo, we used the SmoM2 oncogenic and constitutively active version of Smo to activate the pathway. This form of Smo is localizes to cilia and activates the pathway downstream of Smo regardless of upstream events. To measure pathway activation, we used qRT-PCR to measure *Gli1* expression as this gene is highly upregulated by hedgehog signaling. Wild type mouse embryonic fibroblasts (MEFs) transfected with a SmoM2-mCherry construct showed high levels of *Gli1* expression comparable to what is seen in non-transfected cells treated with the pathway agonist smoothened agonist (SAG) (Fig.1A). In contrast, SmoM2 expression fails to fully activate *Gli1* expression in *Ift27* null MEFs indicating that IFT27 is necessary for signal transduction events downstream of smoothened (Fig. 1A).

**Figure 1.**
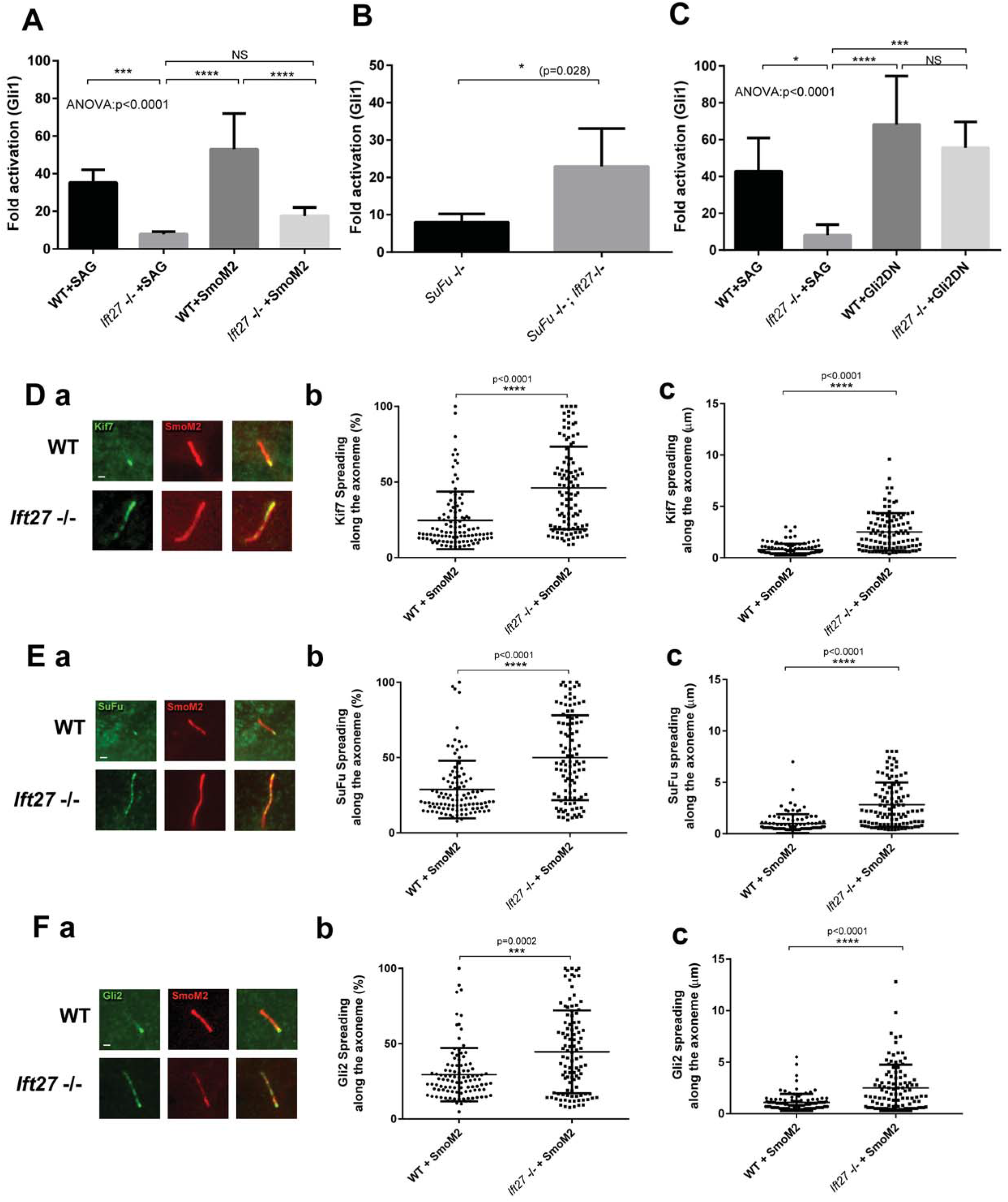
IFT27 is fully integrated in hedgehog signaling pathway. **A.** qRT-PCR quantification of *Gli1* expression in wild type and *Ift27* knockout cells activated by SAG or by the transfection of *SmoM2*. Five independent analysis were performed. **B.** qRT-PCR quantification of *Gli1* expression in *SuFu* knockout cells or *SuFu*, Ift27double knockout cells. Five independent analysis were performed. **C.** qRT-PCR quantification of *Gli1* expression in wild type and *Ift27* knockout cells activated by SAG or by the transfection of constitutively active form of Gli2 (Gli2ΔN). Five independent analysis were performed. **D-F.** Localization of hedgehog effectors in wild type and *Ift27* knockout cells. Immunofluorescence staining of Kif7 (Da), SuFu (Ea) and Gli2 (Fa). Quantification of the spreading pattern presented as percentage of the axoneme covered by the staining (Db, Eb, Fb) or length (Dc, Ec, Fc). In each case, appx. 120 cilia were measured. Data were analyzed using one-way Kruskal-Wallis test (ANOVA) with Turkeys’s multiple comparison test (A and C), Two-tailed Kolmogorov-Smirnov test (B) or Mann-Whitney test (C, Db, Dc, Eb, Ec, Fb, Fc). Error Bars are SD. Bars, 1 micron.

### Ciliary signal transduction effectors are mislocalized after pathway activation

To understand how the loss of IFT27 affects the pathway downstream of Smo, we examined the localization of the ciliary hedgehog effectors Kif7, SuFu and Gli2 in wild type and *Ift27* mutant cells. In wild type cells, these three proteins localize at the ciliary tip and the amount at the tip increases when the pathway is activated by SmoM2 (Fig. 1D-F). SmoM2 expression in *Ift27* mutant cells increases the ciliary tip localization of these proteins but instead of having a tight focus, the proteins are spread back down the cilium (Fig. 1D-F). This suggests that IFT27 plays a role in transport to the tip or in assembly of the ciliary tip compartment.

### IFT27 is necessary for signal transduction downstream of the cilium but not in the nucleus

SuFu, through sequestration of the Gli transcription factors, is a major repressor of the mammalian hedgehog pathway and its loss leads to the activation of hedgehog signaling (Svärd *et al*, 2006; Tukachinsky *et al*, 2010). As the activation of the pathway by loss of SuFu is independent of the primary cilium (Jia *et al*, 2009), we reasoned that a double knock-out of *SuFu* and *Ift27* would allow us to test the involvement of IFT27 in the post-ciliary steps of hedgehog transduction. To do this, we used CRISPR/CAS9 to knock out *Ift27* in *SuFu* null cells (Fig.S1A). If the pathway downstream of SuFu is independent of IF27, then the loss of IFT27 should have no effect on the increase in *Gli1* expression seen in *SuFu* null cells. If IFT27 acts to propagate the signal, then *Gli1* expression should be reduced in the double mutants. Surprisingly, the double knock-out shows a greater activation of the pathway, indicating that IFT27 is a repressor of the pathway at the level of SuFu or downstream (Fig. 1B).

To test the involvement of IFT27 in the activation of target genes in the nucleus, we examined how expression of Gli2ΔN affects gene expression in *Ift27* mutants. This version of Gli2 lacks the N-terminal repressor domain and constitutively activates the pathway (Tanimura *et al*, 1998). No differences in *Gli1* expression were seen between wild type and mutant *Ift27* cells, indicating that IFT27 is dispensable for Gli2 entry in the nucleus and activation of target genes (Fig.1C).

### BBSome and LZTFL1 have different roles in hedgehog signaling

Defects in the BBSome and the BBSome regulator LZTFL1 (Schaefer *et al*, 2014; Seo *et al*, 2011) cause retention of Smo in cilia of non-activated cells like what we observe in *Ift27* mutant cells. Since defects in IFT27 also disrupt the ciliary localization of the BBSome and LZTFL1, we suggested that IFT27 works through the BBSome and LZTFL1 to remove Smo from cilia of non-activated cells (Eguether *et al*, 2014). To understand whether all the hedgehog defects seen in *Ift27* mutants are due to interactions with the BBSome and LZTFL1, we directly compared the phenotypes resulting from mutations in each. Activation of the pathway by SAG or by expression of SmoM2 showed that *Gli1* expression is attenuated in *Lztfl1* mutant cells but the effect is not as large as seen in the *Ift27* mutants. On the other hand, the BBSome mutant, *Bbs2* (Fig.2B) was less affected than either the *Ift27* or *Lztfl1* mutants. While the differences seen in our *Bbs* cells were similar to published work (75% of normal) (Zhang *et al*, 2012), they did not reach significance. These findings suggest that IFT27 is more integrally involved in hedgehog signaling than LZTFL1 or the BBSome.

**Figure 2.**
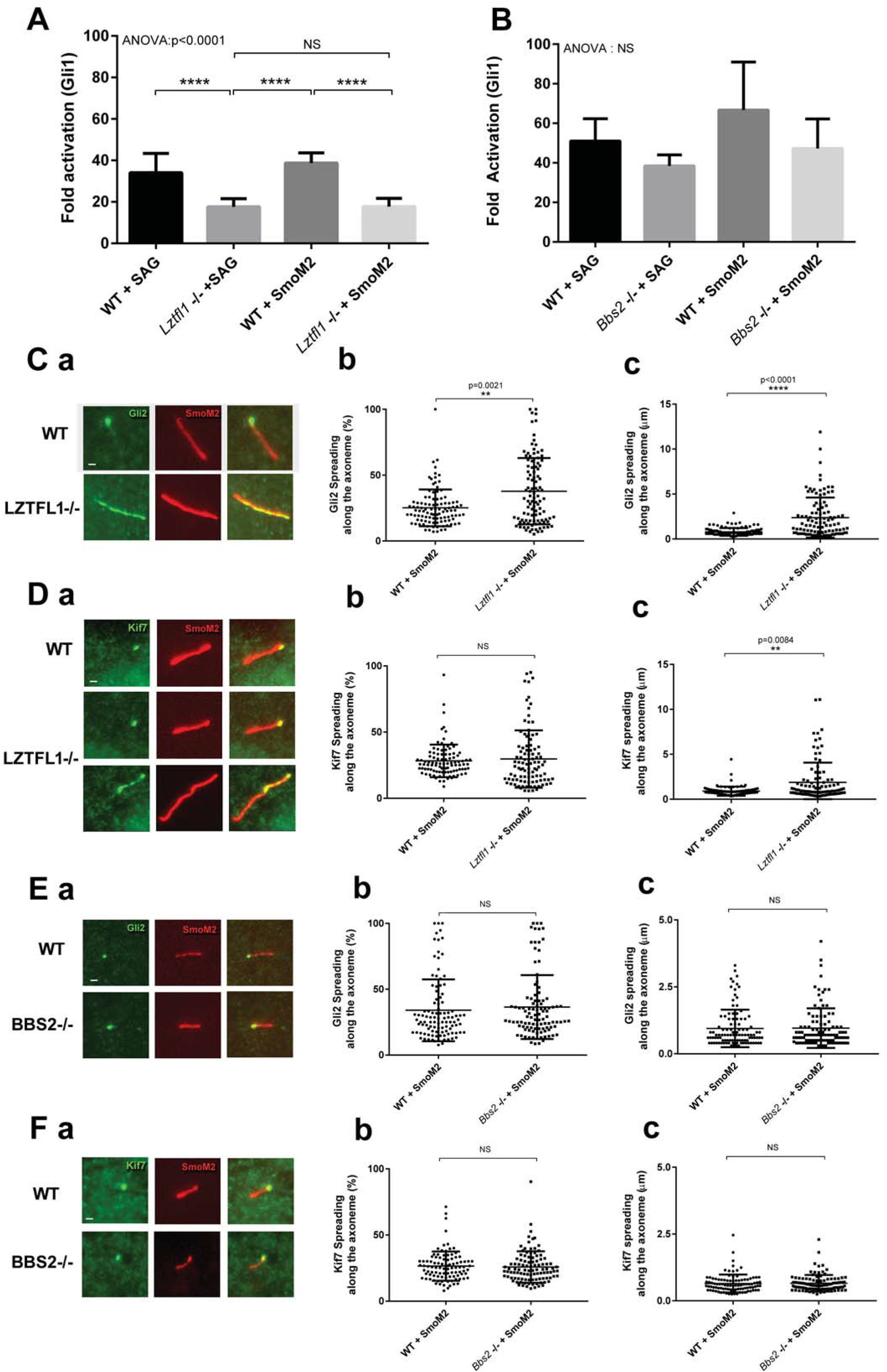
The BBSome and LZTFL1 play different roles in hedgehog signaling. **A.** qRT-PCR quantification of *Gli1* expression in wild type and *Lztfl1* knockout cells activated by SAG or by the transfection of *SmoM2*. **B**. qRT-PCR quantification of *Gli1* expression in wild type and *Bbs2* knockout cells activated by SAG or by the transfection of *SmoM2*. **C-D.** Localization of Hedgehog effectors in wild type and *Lztfl1* knockout cells. Staining of Gli2 (Ca) and Kif7 (Da). Quantification of the spreading pattern in percentage of the axoneme covered by the staining (Cb, Db) and length (Cc, Dc). In each case, appx. 120 cilia were mesured **E-F**. Localization of Hedgehog effectors in wild type and *Bbs2* knockout cells. Staining of Gli2 (Ea) and Kif7 (Fa). Quantification of the spreading pattern in percentage of the axoneme covered by the staining (Eb, Fb) and length (Ec, Fc). In each case, appx. 120 cilia were measured. Data were analyzed using one-way Kruskal-Wallis test (ANOVA) with Turkeys’s multiple comparison test (A and B) or Mann-Whitney test (Cb, Cc, Db, Dc, Eb, Ec, Fb, Fc). Error Bars are SD. Bars, 1 micron.

To more carefully examine the functions of LZTFL1 and the BBSome in hedgehog signal propagation, we expressed SmoM2 in *Lztfl1* and *Bbs2* mutants and quantitated the effects on ciliary Kif7 and Gli2. As observed in *Ift27* mutants, *Lztfl1* mutants showed expansion of the zone of Gli2 and Kif7 label down from the tip of the cilium (Fig.2C,D). Analysis of the results is complicated by the fact that expression of SmoM2 in *Lztfl1* mutant cells increased ciliary length. This caused the percent of the cilium labeled by Kif7 to not be significantly different between mutants and controls even though the zone was physically larger in the mutants (Fig.2Db,c). In *Bbs2* cells, no difference was found between wild type and mutant cells for either Gli2 or Kif7 staining (Fig. 2E,F).

Overall, our qRT-PCR and immunofluorescence experiments show minimal involvement of BBS2 in hedgehog signaling. LZTFL1 is important for hedgehog signaling but is less involved than IFT27. This suggests that IFT27 has BBSome-independent functions and further suggests that LZTFL1 also plays roles independent of the BBSome.

### IFT27 and LZTFL1 cells are defective in Gli2/3 expression and processing

Our experiments expressing SmoM2 in *Ift27* mutants suggests that IFT27 has critical functions downstream of Smo but upstream of events in the nucleus. Possible roles for IFT27 in these steps include regulating the levels of the Gli factors or regulating the conversion of the transcription factors from active to inactive forms. To test these ideas, we examined the levels of Gli2 and Gli3 in wild type, *Ift27^-/-^, Lztfll^-/-^* and *Bbs2^-/-^* cell lines expressing SmoM2 compared to untransfected cells (Fig.3A). All values were normalized to GAPDH and then to either the SmoM2-untransfected counterparts (Fig.3.B,C) or to each of the wild type untreated cell lines (Fig.3D,E).

**Figure 3.**
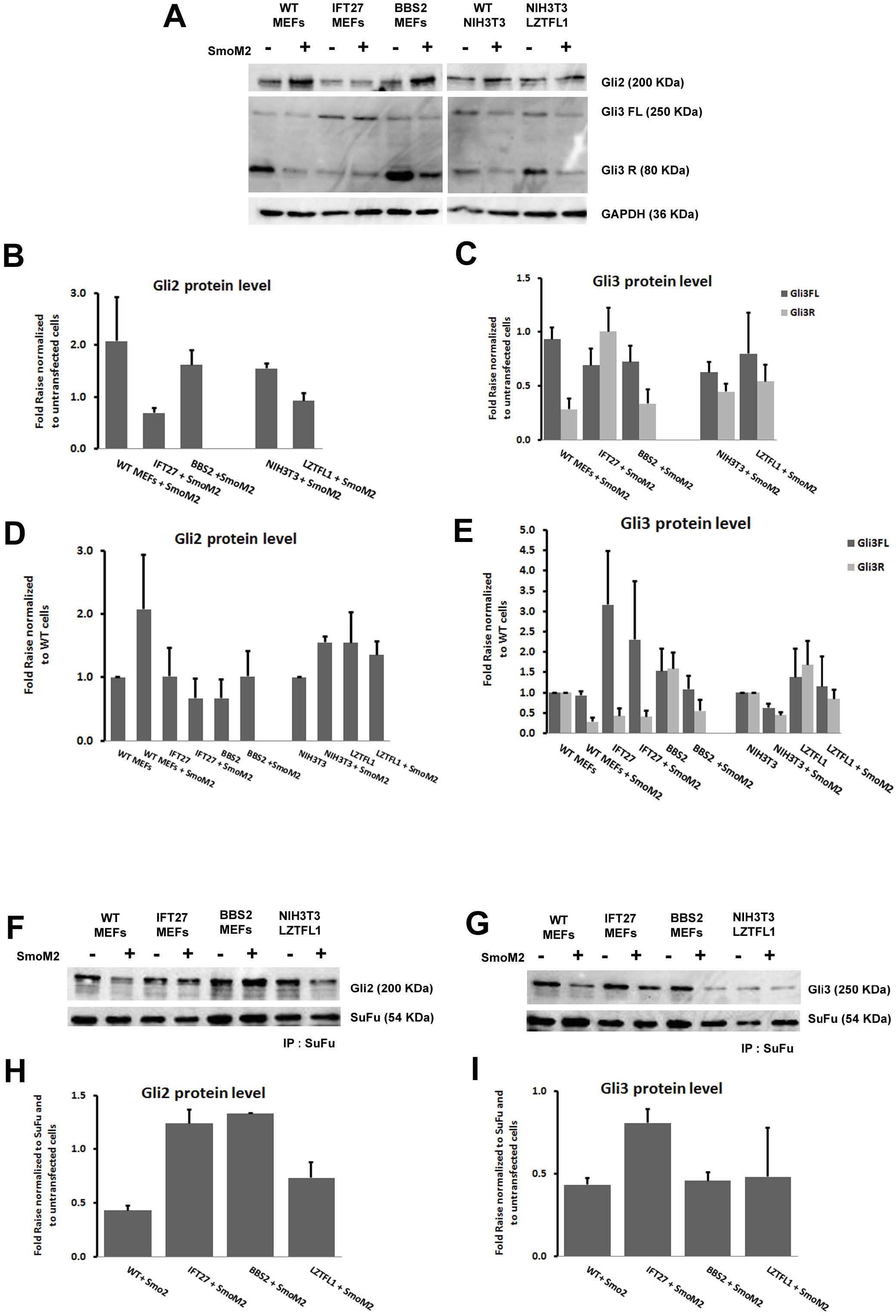
*Ift27* and *Lztfl1* mutant cells are defective in Gli2/3 expression and processing. **A-C.** Immunoblots of Gli2 and Gli3 in wild type, *Ift27, Bbs2* and *Lztfl1* knockout cells transfected or not with SmoM2. GAPDH is the loading control. Quantification of Gli2 (B), Gli3 full length (Gli3FL) and repressor form (Gli3R) (C) expression from three experiments normalized to untransfected cells expression. **D-E.** Quantification of Gli2 (D), Gli3 full length (Gli3FL) and repressor form (Gli3R) (E) expression from three experiments normalized to wild type cells expression. (F) WT, *Ift27* **F.** Pairs of control and SmoM2-transfected cells were lysed, immunoprecipitated with SuFu antibody and analyzed by immunoblotting with Gli2 antibody to evaluate SuFu/Gli2 binding. **G.** Pairs of control and SmoM2-transfected cells were lysed, immunoprecipitated with SuFu antibody and analyzed by immunoblotting with Gli3 antibody to evaluate SuFu/Gli3 binding. **H-I**. Quantification of Gli2 (H) and Gli3 Full Length (I) expression from three experiments normalized to untransfected cells expression. Graphs represent three technical replicates. Error Bars are SD

In wild type cells, the level of Gli2 protein, (the main activator at the transcription level) increases about two fold upon pathway activation in MEFs and about 1.5 fold in NIH 3T3 cells (Fig.3B). *Bbs2* mutants behave like wild type, consistent with our failure to detect significant defects in signaling downstream of Smo in the cells. Interestingly, *Ift27^-/-^* and *Lztfl1^-/-^* cells showed decreases in the level of basal Gli2 and did not show an increase after pathway activation (Fig.3B,D).

In wild type cells, Gli3 exists as a 250kD full length isoform that is thought to have limited ability to activate the pathway and an 80kD isoform with only repressive activity (Fig.3A) (Wang & Li, 2006). In control unstimulated cells, Gli3 is mostly converted to the repressor form, which is destroyed upon pathway activation. *Bbs2^-/-^* and *Lztfl1^-/-^* cells behave much like wild type. However, *Ift27^-/-^* cells have increased levels of full length Gli3 and show little processing to the repressor form (Fig.3C, E). This increased level of full length Gli3 seen in *Ift27* mutants may explain the increased activation seen in *SuFu*^-/-^, *Ift27^-/-^* cells as compared to *SuFu^-/-^* cells.

At the basal state, SuFu binds the full length forms of the Gli transcription factors to stabilize and sequester to prevent activation of gene expression. Upon pathway activation, SuFu releases the Gli transcription factors to allow their activation and participation in transcription (Tukachinsky *et al*, 2010). To understand if the loss of IFT27 altered the interactions between SuFu and Gli2 or Gli3 we precipitated SuFu from controls and cells expressing SmoM2 and quantified the amount of Gli2 and Gli3 that coprecipitates (Fig. 3F-I). As expected, activation of wild type cells reduced the amount of Gli2 and Gli3 that bound SuFu. *Lztfl1* mutants behaved like wild type for both Gli2 and Gli3. *Bbs2* mutants showed normal behavior regarding Gli3 but had abnormally high levels of binding to Gli2 after pathway activation. *Ift27* mutants had abnormally high binding of SuFu to both Gli2 and Gli3 after pathway activation suggesting that IFT27 participates in the release of the Gli transcription factors from SuFu.

### Development of a system to stop IFT after ciliary assembly

In order to study the role of IFT in ciliary signaling without the complication that IFT is required to build the organelle, we developed a rapamycin-dimerization approach to temporally perturb IFT. To do this, we created lines where *Ift* null cells were transfected with IFT-FKBP and FRB- MAO-A expression constructs. In the absence of rapamycin, the FRB-MAO-A fusion protein is sequestered at the mitochondria and the IFT-FKBP participates in IFT to assembly cilia. The addition of rapamycin dimerizes the FKBP and FRB domains, sequestering the IFT protein at the mitochondria and preventing its participation in IFT (Fig.4A).

**Figure 4.**
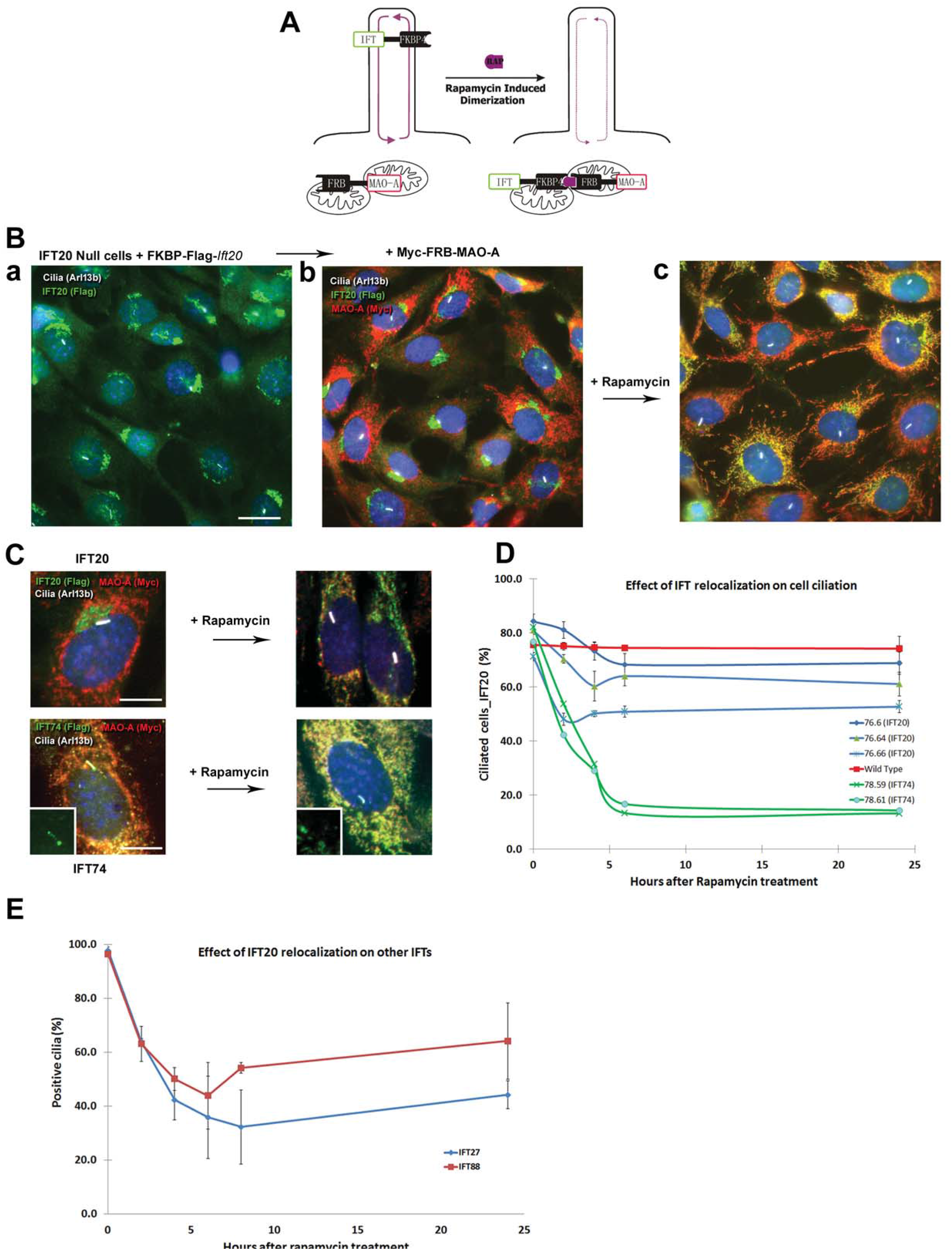
Sequestering IFTs at the mitochondria. **A.** Schematic of the rapamycin-induced sequestration process. The FKBP fragment is fused to mitochondrial membrane protein Monoamine Oxidase A (MAO-A). The FRB fragment is fused to an IFT. Addition of rapamycin traps the IFT fusion protein in the mitochondrial compartment. **B.** *Ift20-null* cells were rescued with an FKBP-Flag-*Ift20* construct (a) and then transfected with an Myc-FRB-*MAO-A* construct (b). Addition of rapamycin relocates IFT20 from the Golgi complex to the mitochondria (c). Bar is 10 microns. **C.** Higher magnification images showing the relocalization of IFT20 and IFT74 by rapamycin treatment. Bar is 5 microns. **D.** Quantification of percent ciliated cells after treatment with rapamycin in control cells, three clones expressing the IFT20-Flag-FKBP construct and two clones expressing the IFT74-Flag-FKBP construct In each case, appx. 120 cells were counted. **E.** Quantification of IFT27 and IFT88 positive cilia after treatment with rapamycin in IFT20 cells (clone TE79.6). In each case, appx. 120 cells were counted. Error Bars are SD

For these experiments, we transfected *Ift20* mutant MEFs and NIH-3T3 *Ift74* mutant CRISPR cells (Fig.S1B) with TE79 (4xFKBP-3xFlag-Mm*Ift20*) and TE78 (Mm*Ift74*-3xFlag-4xFKBP) respectively. Like wild type IFT20, the IFT20-FKBP fusion is localized at the Golgi complex and cilium while IFT74-FKBP is localized to the cilium and around the base of the cilium. Both constructs rescue the ciliation defects present in the parental lines (Fig.4B,C). Uniformly expressing cell lines were obtained by dilution cloning and transfected with TE76 (3xMyc-FRB-*MAO-A*), which expresses a mitochondrial targeted FRB domain (Fig.4B,C). After another round of dilution cloning, cell lines with functional cilia were identified by their ability to activate the hedgehog pathway and accumulate smoothened in cilia after SAG stimulation.

Rapamycin treatment relocalized the IFT74-FKBP and the IFT20-FKBP to the mitochondria in both lines which resulted in a marked depletion of the cilia and Golgi pools of proteins (Fig. 4C).

### Sequestration of IFT20 or IFT74 leads to cilia resorption

In *Chlamydomonas*, temperature sensitive IFT mutants gradually shorten their cilia starting about two hrs after shift to non-permissive temperatures (Huang *et al*, 1977). To understand if mammalian cilia behave similarly, we followed percent ciliation after treatment with rapamycin. Three IFT20 and two IFT74 cell lines were chosen for analysis (Fig.4D). The three IFT20 lines showed a moderate but robust diminution of ciliation ranging from 23% for line 76.66 to 15% for line 76.6. Curiously, ciliary loss plateaued after a few hours and did not continue to decrease. Rapamycin has no effect on the percentage of ciliated wild type cells. For IFT74, deciliation was more extensive and both lines lost approximatively 60% of their cilia by 6 hours of treatment. The difference in efficiency of deciliation between IFT74 and IFT20 may be related to the abundance of the two proteins. The plateau is not fully understood but may be caused by saturation of the FRB binding sites on the mitochondria since rapamycin binding to targets is irreversible. Line 76.6 was chosen for more study as it had robust removal of IFT20 from the Golgi complex and cilium but retained cilia the longest, giving more time for analysis before cilia loss.

If the sequestration of IFT20 or IFT74 at the mitochondria disrupts IFT, we would expect that other IFT proteins would be lost from the cilium. To test this, we quantified the percent cilia positive for IFT88 and IFT27 after rapamycin treatment. At the start of the experiment, nearly all ciliated cells showed the expected staining pattern with IFT27 and IFT88 antibodies. Addition of rapamycin caused a gradual decrease in the percent positive cilia to about 40% of the population before the percent plateaued and started to rebound (Fig.4E).

### Ciliary entry of Smo is independent of IFT

Our previous work on IFT25 and IFT27 implicated IFT in the removal of Smo from cilia. This work clearly showed that IFT25 and IFT27 were not needed for the entry of Smo but other parts of the IFT system could still be involved in Smo transport to cilia. To test this possibility, we asked how perturbing IFT would affect the delivery of Smo to cilia in response to pathway activation. We reasoned that if the IFT system was involved in Smo transport, accumulation of Smo in the cilium upon SAG activation should be blocked or attenuated by addition of rapamycin to our cell system. Cells were treated with SAG for 1 hr and then treated with rapamycin. Cells were fixed at intervals and ciliary Smo quantified. If IFT is required for Smo delivery, the level of ciliary Smo should not increase as fast in the rapamycin-treated cells compared to non-rapamycin treated cells. Quantification showed the same accumulation profile regardless of rapamycin treatment, demonstrating that Smo entry in the cilium is independent of the IFT system (Fig.5A).

**Figure 5.**
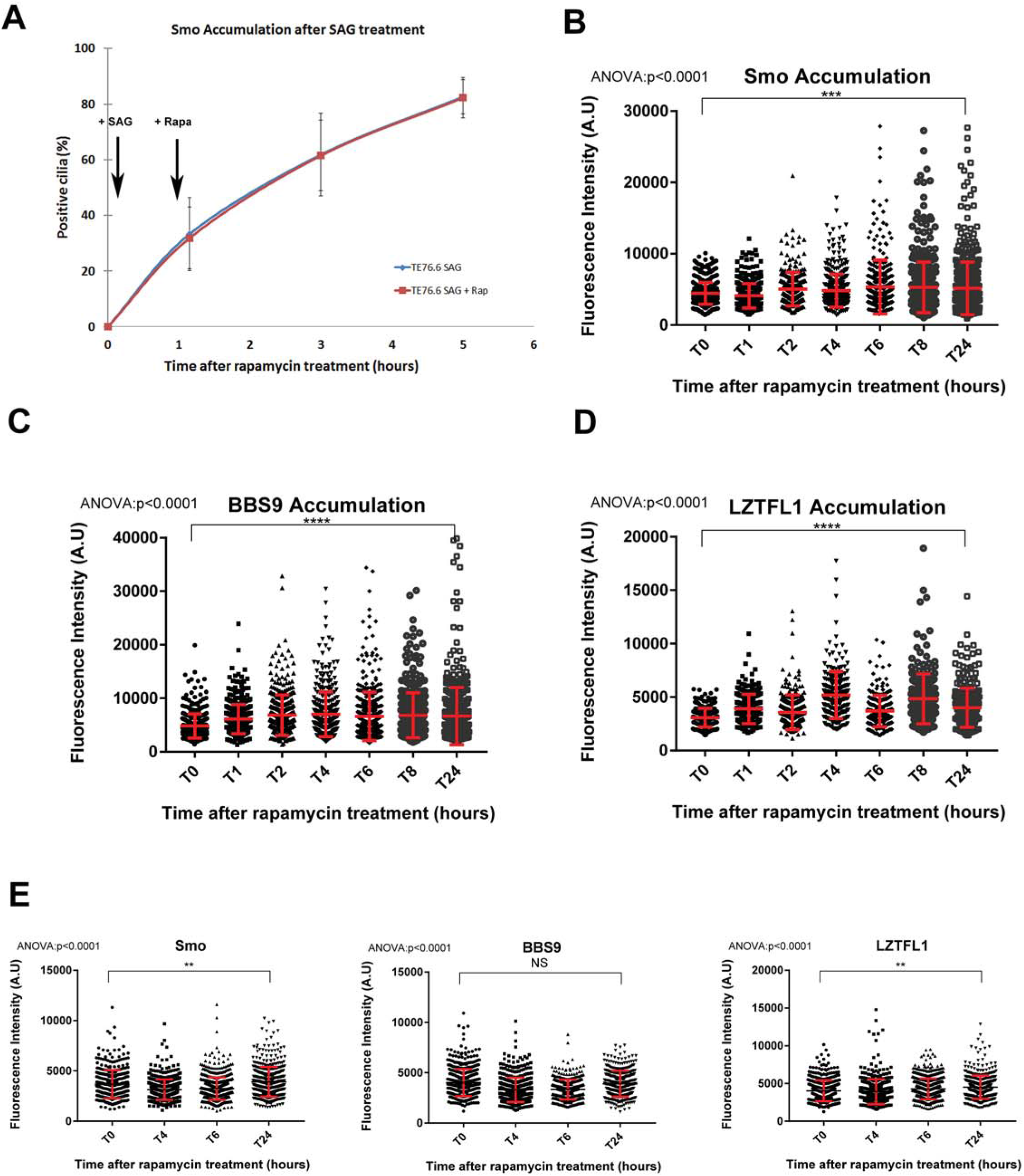
Independent entry of hedgehog effectors in the cilium. **A.** Quantification of Smo positive cilia after activation of the pathway with SAG (added at 5 minutes) and treatment with rapamycin (added at 1h to half of the samples). **B.** Quantification of Smo fluorescence intensity after activation of the pathway under the same experimental paradigm as A. **C.** Quantification of BBS9 fluorescence intensity after activation of the pathway under the same experimental paradigm as A. **D.** Quantification of LZTFL1 fluorescence intensity after activation of the pathway under the same experimental paradigm as A. **E.** Quantification of Smo, BBS9 and LZTFL1 fluorescence intensity over time in rapamycin-treated unstimulated wild type cells. Data were analyzed using one-way ANOVA with Dunett’s multiple comparison test. In each panel, appx. 300 cells were counted. Error Bars are SD

To confirm that Smo entry is IFT independent, we quantified Smo accumulation in cells with IFT impaired by rapamycin but not stimulated with ligand. If Smo delivery is independent of IFT, we expect these cells to accumulate Smo like we observed in cells missing IFT25 or IFT27 (Keady *et al*, 2012; Eguether *et al*, 2014). Over time, Smo increased in cilia on IFT- impaired cells while treatment of wild type cells with rapamycin had no effect on ciliary Smo levels (Fig.5B,E) thus confirming that entry of Smo in the primary cilium is independent of the IFT system.

In addition to accumulating Smo, *Ift27*-mutant cells also accumulated BBS9 and LZTFL1. We assumed these BBS-related components were delivered by other IFT proteins and removed by IFT25/IFT27. If this is true, BBS components should not accumulate in cells with IFT impaired by rapamycin. Surprisingly, the treatment of our cells with rapamycin caused an accumulation of both proteins (Fig.5C, D). Wild type cells treated with rapamycin showed no accumulation of either BBS9 or LZTFL1 in the cilium (Fig.5E). These findings suggest that BBS9 and LZTFL1 entry in the cilium is at least partially independent of the IFT system whereas their removal depends on IFT proteins.

## Discussion

### IFT27 is deeply integrated into the hedgehog pathway

In this work we show that *Ift27* mutant cells have defects in almost every step of the hedgehog signal transduction cascade (Table 1). The defects start at the beginning with abnormal accumulation of Smo in non-activated cilia and a failure to remove Ptch1 after activation. Farther down the pathway, we find defects in the localization of Gli2, Kif7 and SuFu at the ciliary tip. In addition, *Ift27* mutants have increased levels of the full length Gli3 transcription factor, which is not correctly processed to the repressor form in the absence of pathway stimulation. Unexpectedly, double mutant *SuFu^-/-^, Ift27^-/-^* cells showed higher levels of basal *Gli1* expression than single mutant *SuFu^-/-^* cells suggesting that this increased level of full length Gli3 in *Ift27* mutants has the ability to activate *Gli1* expression if it is not repressed by SuFu.

**Table 1.** Summary of hedgehog pathway defects.

|  | *Bbs2*<sup>-/-</sup> | *Lztfl1*<sup>-/-</sup> | *Ift27*<sup>-/-</sup> |
| --- | --- | --- | --- |
| Removal of Ptch1 | not tested | not tested | **impaired** |
| Removal of Smo | **impaired** | **impaired** | **impaired** |
| Kif7 at ciliary tip | normal | **unfocused** | **unfocused** |
| Gli2 at ciliary tip | normal | **unfocused** | **unfocused** |
| SuFu at ciliary tip | not tested | not tested | **unfocused** |
| [Gli3FL] at basal level | normal | normal | **increased** |
| Gli2 release from SuFu | **abnormal** | normal | **abnormal** |
| Gli3 release from SuFu | normal | normal | **abnormal** |

The only step where we do not see an IFT27 effect is the activation of gene expression by truncated Gli2 suggesting that nuclear entry and transcriptional stimulation does not require IFT27. Interestingly IFT25, the obligate binding partner of IFT27, was originally identified as a nuclear protein upregulated in neuroectodermal tumors (Pozsgai *et al*, 2007). These tumors included medulloblastomas, which are known to be driven by upregulated hedgehog signaling (Wu *et al*, 2017), suggesting that there may be undiscovered roles for IFT25 and IFT27 in the nucleus.

### IFT27 has BBSome-dependent and BBSome-independent roles in hedgehog signaling

As *Lztfl1* and *Bbs* mutants share the Smo accumulation phenotype with *Ift27* mutants and LZTFL1 and BBS proteins accumulate in *Ift27^-/-^* cilia, we suggested that IFT27 functions through LZTFL1 and the BBSome. Our current studies show that *Ift27* and *Lztfl1* but not *Bbs2* mutants have defects in the localization of Kif7 and Gli2 at the ciliary tip and *Ift27*-mutants have increased levels of full length Gli3 not seen in *Bbs* or *Lztfl1* mutants (Table 1). The extent of the protein’s involvement in the pathway correlates with severity of the corresponding mouse phenotypes. *Ift27* mutants are most strongly affected with no animals living past the day of birth. About one half of *Lztfl1* mutants die *in utero*. The remaining survivors become obese and develop retinal degeneration (Datta *et al*, 2015; Jiang *et al*, 2016). *Bbs2* mutant mice are initially smaller than littermates and develop obesity, retinal degeneration and kidney cysts with age (Nishimura *et al*, 2004). While numerous hedgehog-related phenotypes are observed in *Ift27* mutants, no hedgehog phenotypes such as polydactyly, cleft palate, holoprosencephaly, cardiac malformations or lung isomerisms were reported in the *Lztfl1* or *Bbs2* mutant mice. These data indicate that both IFT27 and LZTFL1 have BBSome-independent functions in hedgehog signaling.

### Development of a system to uncouple IFT role in ciliogenesis from signaling

Temperature sensitive alleles of IFT components are powerful tools for uncovering the role of IFT in fully formed cilia. The difficulty in obtaining temperature sensitive alleles in mice drove us to develop an alternative approach based on rapamycin-induced in-cell dimerization to perturb IFT in mature cilia. Previously we had shown that Smo entry was independent of IFT25 and IFT27. Our new system demonstrated that SAG-stimulated Smo entry was not dependent on IFT in general. Also, Smo accumulated in non-stimulated cilia when IFT was perturbed supporting a role for IFT in removal of Smo but not in the entry. This data is consistent with published work showing that Smo accumulates in cilia on cells with defective IFT dynein (Ocbina & Anderson, 2008) and with recent work from our group showing that Smo entry into cilia does not utilize IFT20 (Monis *et al*, 2017). Smo may be an unusual membrane protein as it is thought to enter the ciliary compartment from a plasma membrane pool rather than from an internal membrane pool (Milenkovic *et al*, 2009). Our newly developed tool will allow us to address the role of IFT in the delivery of other membrane proteins.

### Entry of the BBSome may be IFT-independent

In *C. elegans* and *Chlamydomonas* the BBSome is transported through cilia by IFT (Ou *et al*, 2005; Lechtreck *et al*, 2013) and it is expected to be carried into cilia by the IFT kinesin motors as part of a large complex with IFT-A and IFT-B. Unexpectedly we found that LZTFL1 and BBS9 accumulate in cilia when we perturb IFT. Similar observations were previously reported in *Chlamydomonas* where BBS4 levels initially dropped after IFT was stopped in a temperature sensitive allele. However, once the IFT proteins were depleted from the cilia, BBS4 started to accumulate. The authors attributed the increase in ciliary BBS4 to increased numbers of cilia being loaded at later time points due to ciliary shortening after temperature shift (Lechtreck *et al*, 2013) but in light of our observations, this interpretation should be reexamined. The experimental approaches in the two experiments were different but the similar observations indicate that BBSome entry into cilia is at least partially independent of IFT. One possibility is that the BBSome can independently engage kinesin-2 for entry. However, BBS4 accumulated in *Chlamydomonas* cilia with kinesin-2 mutations suggesting that a different motor is responsible for initial delivery.

## Materials and Methods

### DNA Constructs

**SmoM2-mCherry**: SmoM2 in pHAGE_DN_CMV vector (gift from D. Nedelcu and A. Salic). TE72 (mNeon-Gli2ΔN): MmGli2 amplified from AA325 to the end and inserted in TE44 vector (mNeonGreen_pHAGE_DN_CMV_Nuc::eGFP) mimicking human Gli2ΔN truncation (Roessler *et al*, 2005).

**TE85** (IFT27 CRIPSR gRNA): IFT27 guide RNAs oligos designed in (Liew *et al*, 2014) were annealed together and inserted in LentiCRISPRv2 plasmid (Gift from Fen Zhang (Sanjana *et al*, 2014) (Addgene Plasmid #52961)).

**TE32** (CRISPR pBSKgRNA with IFT74 oligo #5): Candidate sgRNAs were identified by searching for G(N)20GG motifs 300 bases upstream and 100 bases downstream of the targeting sequence that conform with the nucleotide requirements for U6 Pol III transcription and the spCas9 PAM recognition element (NGG) (Jinek et al., 2012; Mali et al., 2013) using Web-based software ZiFiTtargeter 4.2 (Sander et al., 2010). Sequences generated were aligned to mouse genome using nBLAST to search for potential off-target sites. Pairs of oligonucleotides were subsequently annealed together and cloned into pBSK-gRNA (Gift of R. Maehr). IFT74 target sequence is GTTCTCGTGGTGGTCCCTTA.

**TE76** (3xMyc-FRB-MAO-A in pHAGE-DN-CMV-nuc::eGFP): The FKBP-Rapamycin binding domain (FRB) fused to Mm MAO-A was PCR amplified from a CFP-FRB-MAO-A construct (Gift of T. Inoue) with the 3xMyc tag in the 5’ primer and cloned into the lentiviral vector pHAGE-DN-CMV-nuc::eGFP.

**TE78** (MmIFT74-3xFlag–4xFKBP in pHAGE-puro): Four tandem repeats (4xFKBP) of the FK506 Binding Protein were PCR amplified from a vector containing IFT20-4xFKBP12 (Gift of T. Inoue). IFT74-3xFlag was amplified from TE60. Both fragments were inserted in the lentiviral vector pHAGE-puro with the Gibson Assembly system (New England Biolabs).

**TE79** (4xFKBP-3xFlag-MmIFT20 in pHAGE-puro): Four tandem repeats (4xFKBP) of the FK506 Binding Protein and IFT20-3xFlag fragments were PCR amplified from a vector containing IFT20-4xFKBP12 (Gift of T. Inoue). Both fragments were inserted in the lentiviral vector pHAGE-puro with the Gibson Assembly system (New England Biolabs).

### Cell culture

*Ift27* and *Lztfl1* mutant cell lines are described elsewhere (Eguether *et al*, 2014). *Bbs2* mutant MEFs were isolated from E13.5 *Bbs2* mutant embryos (Nishimura *et al*, 2004) and immortalized with the large T antigen from SV40 virus. The *SuFu* mutant cell line was a gift from A. Salic. All DNA constructs were transfected into the cells via lentiviral infection followed by drug selection to create stable cell lines.

NIH3T3 fibroblasts, MEFs and their derivatives were grown at 37°C in 5% CO_2_ in Dulbecco’s modified Eagle’s medium (DMEM; Gibco) with 10% FBS and 1% Penicillin-Streptomycin (Gibco). For SAG experiments, cells were plated at near confluent densities and serum starved (same culture medium described above but with 0.25% FBS) for 48 hours prior to treatment to allow ciliation. SAG (Calbiochem) was used at 400nM.

To generate IFT74 rapamycin dimerization cells (NIH3T3 + TE32cl.33 + TE78 + TE76), NIH 3T3 were electroporated with TE32 and the Cas9-puro vector to knock out IFT74, selected with puromycin and dilution cloned. After western blot screening for loss of IFT74, clone 33 was selected and transfected with TE78. After blasticidin selection, cells were dilution cloned, screened for rescue of primary cilium formation and ability to activate the hedgehog pathway under SAG stimulation. Clone 71 was selected and transfected with TE76.

To generate IFT20 rapamycin dimerization cells (14176.1T MEFs + TE79 + TE76), MEFs from *Ift20* mutant embryos (E9.5) were immortalized with the Large T antigen from SV40. Immortalized cells were transfected withTE79, blasticidin selected, dilution cloned and screened for rescue of primary cilium formation and ability to activate the hedgehog pathway upon SAG stimulation. Clone 16 was selected and transfected with TE76.

Cell lines were generated from mouse models or edited by CRISPR and were authenticated by genotyping loci of interest to the study. Presence of mycoplasma is monitored by DAPI stain.

### Lentivirus Production

Lentiviral packaged pHAGE-derived plasmids (Wilson *et al*, 2008) were used for transfection. These vectors are packaged by a third generation system comprising four distinct packaging vectors (Tat, Rev, Gag/Pol, VSV-g) using HEK 293T cells as the host. DNA (Backbone: 5μg; Tat: 0.5 μg; Rev: 0.5 μg; Gag/Pol: 0.5 μg; VSV-g: 1 μg) was delivered to the HEK cells using Effectene (Qiagen) or as calcium phosphate precipitates. After 48 hours, supernatant was harvested, filtered through a 0.45 mm filter and precipitated with Lenti-x concentrator (Clontech). Precipitated viral particles were resuspended in DMEM with polybrene (5mg/mL) and added to 10-20% confluent cells. After 24 hrs, cells were selected with antibiotic.

### Protein and mRNA Analysis

For western blots, MEFs were lysed directly into denaturing gel loading buffer (Tris-HCl 125mM pH6.8, Glycerol 20% v/v, SDS 4% v/v, β-Mercaptoethanol 10% v/v, bromophenol blue). Western blots were developed by chemiluminescence (Super Signal West Dura, Pierce Thermo) and imaged using a LAS-3000 imaging system (Fujifilm).

For immunoprecipitations, cells were serum starved for 48 hours and proteins were extracted with lysis buffer (20 mM Hepes pH 7.5, 50 mM KCl, 1mM MgCl2) with 0.5% digitonin and protease inhibitor (Complete EDTA-Free, Roche). Insoluble components were removed by centrifugation at 20000g. Primary antibodies pre-adsorbed to protein G-sepharose beads (GE Healthcare) were added to the cell extract and the mixture incubated for 2 hrs at 4°C. After centrifugation, beads were washed with lysis buffer supplemented with 0.1% digitonin before elution in denaturing gel loading buffer for SDS-PAGE electrophoresis and Western blotting analysis.

Isolation of mRNA and quantitative mRNA analysis was performed as previously described (Jonassen et al., 2008) using the described primers (Table 2).

**Table 2.**
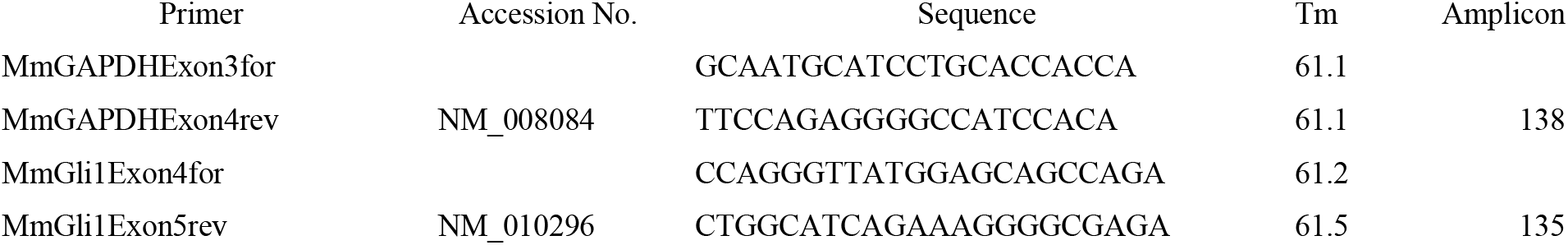
Quantitative RT-PCR primers.

### In Cell Rapamycin Dimerization

IFT20 and IFT74 engineered cells were plated on glass coverslips, grown to confluence and serum starved for 48 hours with DMEM medium containing 0.25% FBS in order to allow cilia growth. To trigger dimerization of the TE76 and TE78/79 constructs, cells were treated with rapamycin (100nM in low-serum medium) and coverslips were then fixed at desired time points.

To study hedgehog signaling in the context of IFT20 removal, cells were plated on glass coverslips, grown to confluence and serum starved for 48 hours with DMEM medium containing 0.25% FBS in order to allow cilia growth. Cells were treated with SAG for one hour to activate the hedgehog pathway. To trigger dimerization of the two constructs, rapamycin was added at a final concentration of 100nM. Coverslips were fixed at various time points.

### Immunofluorescence

Cells were fixed with 2% paraformaldehyde for 15 min, permeabilized with 0.1% Triton-X-100 for 5 min and treated with 0.05% SDS for 5 min to retrieve antigens. The primary antibodies are described (Table 3).

**Table 3.** Antibodies.

| Primary antibody | Clone name | Concentration | Supplier |
| --- | --- | --- | --- |
| Acetylated Tubulin | 6-11B-1 | 1:10000 | Sigma |
| Flag | F1804 | 1:1000 | Sigma |
| Arl13b |  | 1:1000 | Davis/NIH NeuroMab |
| Kif7 |  | 1:1000 | Gift from K.V. Anderson |
| SuFu |  | 1:500 (IF) | Gift from J. Eggenschwiler |
| SuFu | C81H7 | 1:50 (IP) | Cell Signaling |
| Gli2 |  | 1:1000 | Gift from J. Eggenschwiler |
| Gli3 | AF3690 | 1:200 | R and D systems |
| Smoothened | E5 | 1:100 | Santa Cruz |
| IFT27 |  | 1:250 | Lab Made |
| IFT88 |  | 1:250 | Lab Made |

Antibody verification was done by western blot and immunofluorescence. Kif7, SuFu, Gli2, Gli3 and Smo antibodies recognized expected size proteins on western blot and their staining of cilia matched previously published patterns in unstimulated and hedgehog-activated cells. IFT27 and IFT88 antibodies recognized appropriate sized bands on western blots of wild type cells but not from corresponding mutants. By immunofluorescence IFT88 and IFT27 antibodies labeled the mother centriole or basal body in wild type cells but not in corresponding mutants.

### Quantification of fluorescence in cilia

Cilia analysis was performed using an in-house developed ImageJ macro-toolset (Rasband, W.S., ImageJ, U. S. National Institutes of Health, Bethesda, Maryland, USA, https://imagej.nih.gov/ij/, 1997-2016.). Briefly, sets of images are imported into the software and cilia are pre-processed (median filtering and background subtraction), then detected using the “Analyze particle function”. Each cilium is extracted using a 64x64 pixels bounding box. Deconvolution is performed on both the structural marker channel and the signal channel with the DeconvolutionLab plugin (Sage *et al*, 2016), using synthetic PSF generated by the Diffraction PSF 3D plugin (<u>http://www.optinav.info/Diffraction-PSF-3D.htm</u>). For each structure, the following parameters are extracted: total volume, average intensity, Feret length, length of the skeleton, ratio of both the Feret and the skeleton length (an indicator of cilium bending).

### Statistics

Data groups were compared using non parametric Mann-Whitney or Kolmogorov-Smirnov tests (two groups) or one-way Kruskall-Wallis test (more than two groups) followed by Turkey’s or Dunett’s multiple comparison tests, using GraphPad Prism 7 software. Differences between groups were considered statistically significant if p < 0.05. Statistical significance is denoted with asterisks (*p=0.01 – 0.05; **p=0.001-0.01; ***p < 0.001). Error bars are all S.D. Center values are all averages.

## Acknowledgements

We thank our colleagues for sharing reagents described in the materials and methods. This work was supported by funding from the National Institutes of Health GM060992 and DK103632 to GJP. The Bordeaux Imaging Center is a service unit of the CNRS-INSERM and Bordeaux University, member of the national infrastructure France BioImaging supported by the French National Research Agency (ANR-10-INBS-04).

---

**Figure S1.**
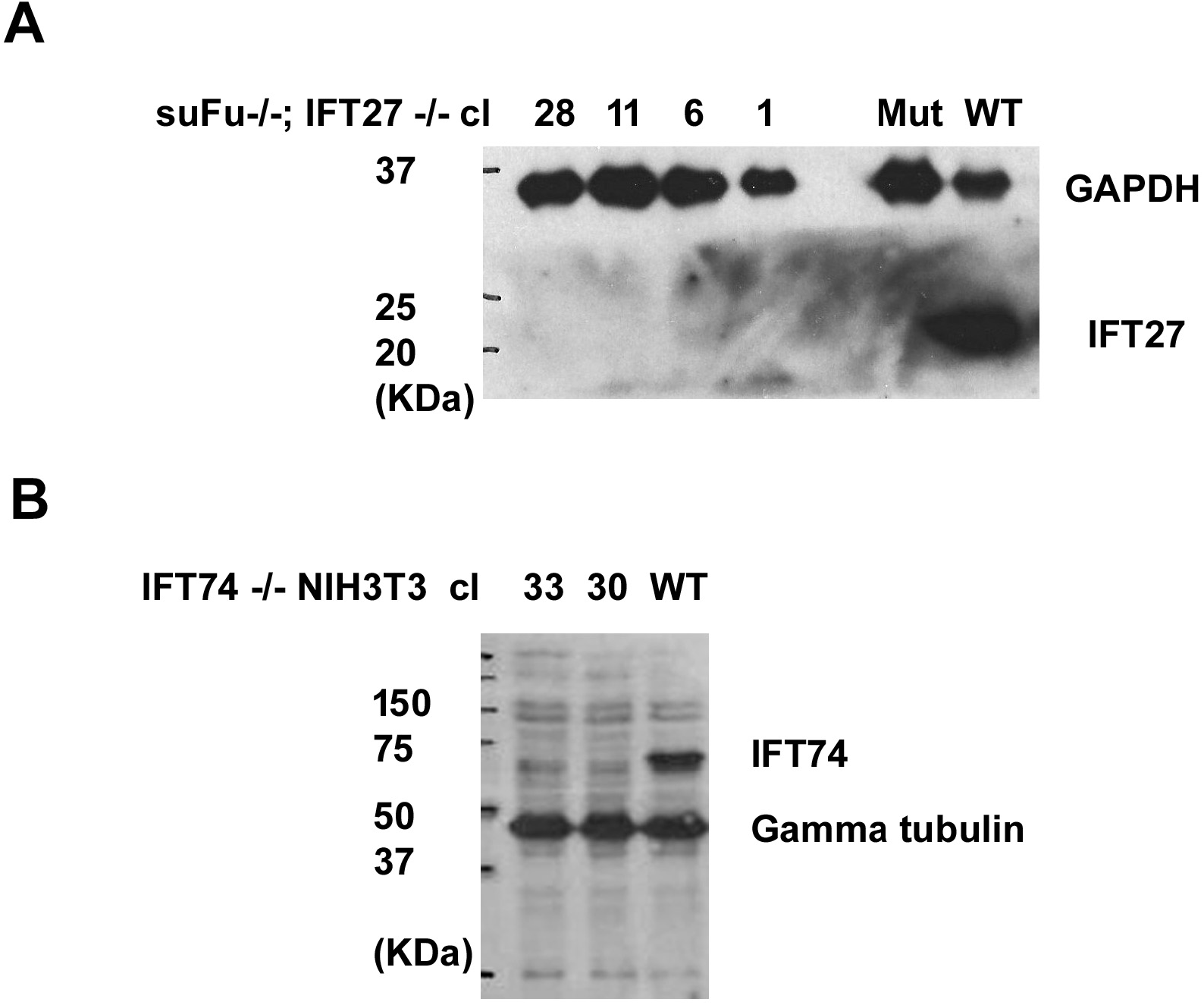
Characterization of CRISPR Knock Out cells. **A.** Immunoblot showing that IFT27 is not expressed in CRISPR *SuFu^-/-^, Ift27^-/-^* lines (28, 11, 6, 1). *Ift27^-/-^* (Mut) and wild type (WT) MEFs are included as controls. GAPDH was used as a loading control. **B.** Immunoblot showing that IFT74 is not expressed in CRISPR *Ift74^-/-^* lines (33, 30) but is present in the parental NIH 3T3 (WT) line. Gamma tubulin was used as a loading control. Error Bars are SD

